# Inferring demographic history using two-locus statistics

**DOI:** 10.1101/108688

**Authors:** Aaron P. Ragsdale, Ryan N. Gutenkunst

## Abstract

Population demographic history may be learned from contemporary genetic variation data. Methods based on aggregating the statistics of many single loci into an allele frequency spectrum (AFS) have proven powerful, but such methods ignore potentially informative patterns of linkage disequilibrium (LD) between neighboring loci. To leverage such patterns, we developed a composite-likelihood framework for inferring demographic history from aggregated statistics of pairs of loci. Using this framework, we show that two-locus statistics are indeed more sensitive to demographic history than single-locus statistics such as the AFS. In particular, two-locus statistics escape the notorious confounding of depth and duration of a bottleneck, and they provide a means to estimate effective population size based on the recombination rather than mutation rate. We applied our approach to a Zambian population of *Drosophila melanogaster*. Notably, using both single– and two-locus statistics, we found substantially lower estimates of effective population size than previous works. Together, our results demonstrate the broad potential for two-locus statistics to enable powerful population genetic inference.

## Introduction

Patterns of genetic variation within a population are shaped by the evolutionary and demographic history of that population, so observed variation encodes information about that history. Knowing population demographic history serves as an important control for learning about natural selection (Bustamante et al., 2001; Boyko et al., 2008) and understanding the relative efficacy of selection as populations change in size (Lohmueller et al., 2008; Henn et al., 2016). One particularly informative statistic used to summarize genetic polymorphism data is the allele frequency spectrum (AFS), which stores the distribution of observed single-locus allele frequencies from a sample of the population. The shape of the AFS is sensitive to demographic history, and fitting the expected AFS under parameterized demographic models to the observed AFS is a powerful approach for learning about demographic history (Marth et al., 2004; Williamson et al., 2005; Gutenkunst et al., 2009; Kamm et al., 2016b).

For unlinked loci, the AFS is a sufficient statistic of the data and completely describes observed patterns of variation (Lohmueller et al., 2009). The expected sample frequency spectrum under arbitrary single– or multi-population histories can be efficiently calculated with either coalescent (Kingman, 1982; Tajima, 1983) or diffusion (Kimura, 1964; Williamson et al., 2005; Gutenkunst et al., 2009) approaches. Poisson random field theory (Sawyer and Hartl, 1992) can then be used to calculate the likelihood of the data given model parameters. A key assumption of the Poisson random field framework is that of independence between segregating loci, so that allele frequency trajectories are uncorrelated. However, neighboring loci are physically linked on the chromosome, and their allele frequencies are thus correlated. Recombination serves to reduce this correlation, with a higher rate of recombination between two loci more rapidly breaking down that association. For any two linked SNPs, their linkage disequilibrium is a measure of their non-independence. Furthermore, as with allele frequencies, patterns of linkage disequilibrium are shaped by historical demographic events such as bottlenecks, growth, and admixture, and therefore they are also informative about history (Pritchard and Przeworski, 2001).

For linked sites the distribution of linkage disequilibrium carries additional information to the allele frequency spectrum about past demography (Myers et al., 2008), and the joint distribution of allele frequencies and linkage disequilibrium between pairs of SNPs should afford greater power for demographic inferences than those based on allele frequencies alone. Characterizing two-locus allele frequency dynamics and calculating their sampling probabilities has attracted a large body of work. Kimura considered the case of genetic drift at multi-allelic loci using a diffusion approximation, and he calculated the time to fixation for one of the alleles when more than two alleles are present (Kimura, 1955). This approach was expanded over the following decade to explicitly consider the two-locus setting with two alleles at each locus (Kimura, 1963; Hill and Robertson, 1966; Karlin and McGregor, 1968; Ohta and Kimura, 1969; Watterson, 1970). These studies were generally interested in the probability and rates of fixation under arbitrary recombination between the two loci and in characterizing the expectation and variance of linkage disequilibrium.

More recently, sampling probabilities for two neutral linked loci were directly calculated under equilibrium demography (Golding, 1984; Hudson, 1985; Ethier and Griffiths, 1990), often using the recursion approach due to Golding (1984). Hudson (2001) extended these results to generate those sampling probabilities with knowledge of the ancestral state and proposed a composite likelihood approach for fine-scale estimation of recombination rates across the genome, which has been implemented to infer recombination maps and identify hotspots in human and *Drosophila* populations (McVean et al., 2004; Auton and McVean, 2007; Chan et al., 2012). Xie (2011) used a diffusion approach to calculate the sample frequency spectrum for two completely linked loci under neutrality or equal levels of selection, while Ferretti et al. (2016) recently used a coalescent approach to calculate the expected frequency spectrum for two completely linked neutral loci, and neutral sampling probabilities were developed under the coalescent with recombination for moderate to large recombination rates and constant population size (Jenkins and Song, 2009, 2010, 2012; Bhaskar and Song, 2012). Recently, Kamm et al. (2016a) developed a coalescent approach to generate two-locus sampling probabilities under arbitrary demography and recombination and found that accounting for demographic history improves accuracy in composite likelihood approaches for estimating fine-scale recombination rates.

Here, we characterize the increase in power of demographic inference from using two-locus allele frequency statistics versus using the single-locus AFS. In particular, the depth and duration of a bottleneck are confounded when using the AFS, but we show they can be independently inferred using two-locus statistics. To enable our analyses, we developed a numerical solution to the diffusion approximation for two-locus allele frequencies with arbitrary recombination. We packaged this method in a two-locus composite likelihood framework that can be used to infer single-population demographic histories. Moreover, this framework allows for an estimate of the effective population size based on recombination that is independent from estimates based on levels of diversity. Using this approach, we inferred demographic history for a highly studied Zambian *Drosophila melanogaster* population, finding a smaller effective population size than previous analyses *N_e_* ∼ 1.5 − 3 × 10^5^ and a demographic history of recent modest growth and no severe bottlenecks.

## Theory and Methods

### A discrete two-locus model with influx of new mutations

We used a diffusion approximation to a two-locus model that allows for two alleles at each locus, which are separated by recombination fraction *r* (Karlin and McGregor, 1968; Watterson, 1970). We allow the left locus to carry alleles *A* and *a*, while the right locus permits alleles *B* and *b*. Then four haplotypes are possible, *AB*, *Ab*, *aB*, and *ab*, with frequencies *n_AB_*, *n_Ab_*, *n_AB_* and *n_ab_* that sum to 2*N* (Fig. 1A). Frequencies in the subsequent generation are found by considering the random pairing of haplotypes and the probability of a given pairing passing on each type to their offspring. These probabilities depend on current haplotype frequencies and the recombination rate and are described in Table 1 of Watterson (1970). For example, a parent carrying haplotypes *AB/Ab* will pass on *AB* with probability 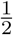 and *Ab* with probability 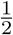, even with recombination. On the other hand, a parent with *AB/ab* will pass on *AB* or *ab* each with probability 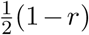 and *Ab* or *aB* each with probability 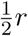. The numbers 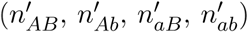 of each haplotype in the next generation are then pulled from the multinomial distribution for sampling 2*N* haplotypes with probabilities found by considering random pairing of haplotypes and recombination.

**Figure 1:**
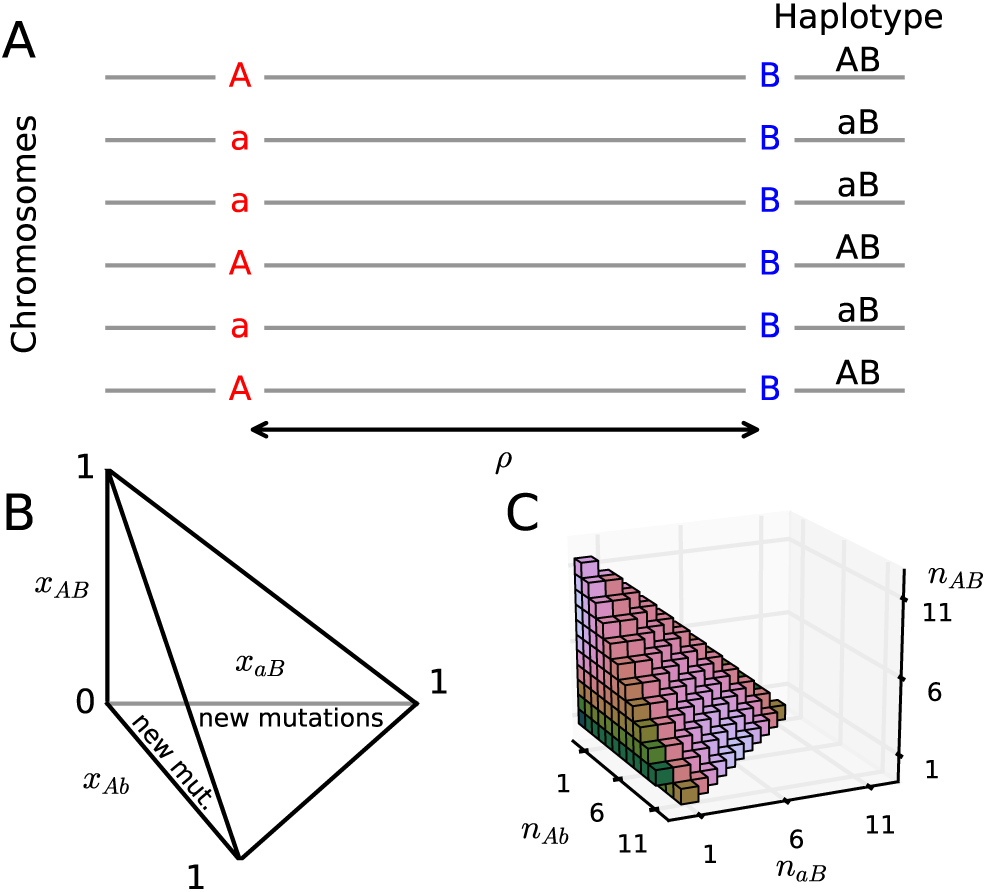
Two-locus model and frequency spectrum. (A) Two loci with two alleles each are separated by recombination distance *ρ* = 4*N_e_r*. Four haplotypes are possible, and we track the frequencies of the three derived haplotypes. (B) Frequencies change within a tetrahedral domain, with corners of the domain corresponding to one of the four haplotypes fixed in the population. New two-locus pairs occur when a new mutation *A* occurs against the *B/b* background, or when *B* occurs against the *A/a* background, so we inject density along the *Ab* or *aB* axes proportional to the background one-locus allele frequencies. (C) A sample two-locus haplotype frequency spectrum for a sample size of *n* = 12.

**Table 1:** Point estimates from fits to *Drosophila* data. Reported log-likelihoods (*LL*) are for two-locus data using the demographic history parameters from each fit. 95% con dence intervals are given in Table S1.

| Data statistics (Model) | $\nu_1$ | $\nu_2$ | $T_1$ | $T_2$ | $p_{\text{mis}}$ | $N_e$ | $LL$ |
| --- | --- | --- | --- | --- | --- | --- | --- |
| One-locus (2-epoch) | 4.23 |  | 0.329 |  | 0.0476 | 302,900 | −1068200 |
| One-locus (3-epoch) | 2.35 | 10.7 | 0.388 | 0.0938 | 0.0496 | 291,500 | −1404200 |
| Two-locus (fix $N_e$, 2-epoch) | 3.83 | | 0.371 | | 0.0449 | $3 \times 10^5$ | −1025600 |
| Two-locus (fix $N_e$, 3-epoch) | 34.3 | 1.69 | 0.220 | 0.053 | 0.0434 | $3 \times 10^5$ | −844200 |
| Two-locus (var. $N_e$, 2-epoch) | 4.02 | | 0.379 | | 0.0456 | 179,900 | −851700 |
| Two-locus (var. $N_e$, 3-epoch) | 1.53 | 4.58 | 0.352 | 0.286 | 0.0473 | 170,000 | −825600 |

New two-locus pairings, with two alleles segregating at both sites, arise when a new mutation occurs at one unmutated locus when the other locus is already polymorphic. Suppose, without loss of generality, that the right locus is already polymorphic, with derived allele *B* at frequency *x_B_* = *n_B_*/2*N*, and ancestral allele *b* at frequency *x_b_* = 1 − *x_B_*. Then a new *A* mutation at the left locus begins at frequency *x_A_* = 1/2*N* and occurs on the *B* haplotype with probability *x_B_* or on the b haplotype with probability *x_b_*. Two-locus frequencies then evolve under the multinomial process described above until one or both loci are fixed for either the ancestral or derived allele, at which point we stop tracking that two-locus pair. The frequencies *x_B_* are drawn from the population distribution of one-locus frequencies *f*(*x*), which can be approximated using diffusion theory (Kimura, 1964). Thus, new independent two-locus pairs enter the population with frequencies (*x_AB_*; *x_Ab_*, *x_aB_*) = (1/2*N*, 0, *x_B_* − 1/2*N*) with rate proportional to *x_B_f(x_B_)* and (0, 1/2*N*, *x_B_*) with rate proportional to (1 − *x_B_*)*f*(*x_B_*).

The density *ϕ*(*x*_1_, *x*_2_, *x*_3_) of two-locus haplotype frequencies, where *x*_1_, *x*_2_ and *x*_3_ are the relative frequencies of haplotypes *AB*, *Ab* and *aB*, respectively (Figure 1B), can be approximated using diffusion theory, as described in the next section. The two-locus haplotype frequency spectrum stores the counts of derived haplotypes in a sample, where one or both loci carry the derived allele. To obtain the two-locus spectrum *F* for *n* samples from the density function *ϕ* (Fig. 1C), we sample against the multinomial sampling distribution:

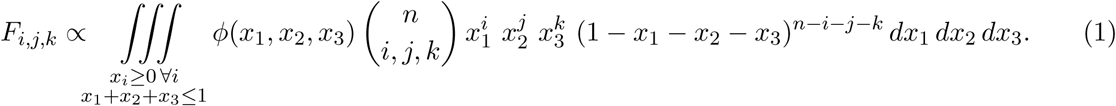

Here, 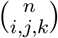 is the multinomial coefficient, defined as 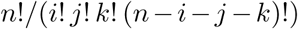. Because we assume that two-locus pairs are independent realizations of this process, Poisson random field theory tells us that if we observe data *D*(*i, j, k*), each entry in the observed two-locus spectrum is a Poisson random variable with mean *F*(*i, j, k*). This allows the application of likelihood theory to compare observed data to model expectations.

### The two-locus diffusion approximation

We solved the multiallelic diffusion equation for *ϕ* to obtain the expected sample two-locus spectrum. Measuring time *τ* in units of 2*N_a_* generations, where *N_a_* is the ancestral reference population size, the forward diffusion equation describes the evolution of the probability density of two-locus frequencies and is written as

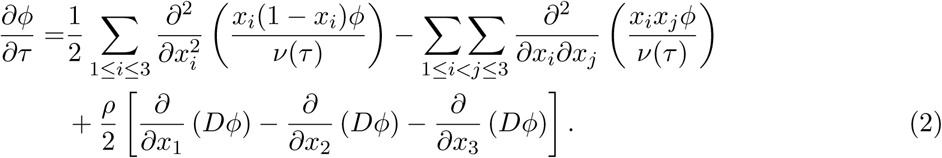

Here, *D* = *x*_1_(1 − *x*_1_ − *x*_2_ − *x*_3_) − *x*_2_*x*_3_ is the linkage disequilibrium, given haplotype frequencies (*x*_1_; *x*_2_; *x*_3_), and 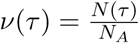 is a function for the relative population size to the ancestral population size at time. The population scaled recombination rate between the *A/a* and *B/b* loci is *ρ* = 4*N_A_r*, where *r* is the recombination rate per generation per meiotic event. The action of recombination is readily interpretable in the diffusion equation; recombination acts directionally on the haplotype frequencies *x_i_*, pushing them toward linkage equilibrium (*D* = 0) at a rate directly proportional to the recombination rate *ρ*.

The domain of the two-locus diffusion equation is the tetrahedron with 0 ≤ *x_i_* ≤ 1 for *i* = 1, 2, 3, and 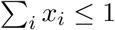(Fig. 1B). If the recombination rate *ρ* = 0 and there is no recurrent mutation, then all boundary surfaces of the domain are absorbing, so if one of the haplotypes is lost from the population it remains lost. However, with *ρ* > 0, the boundary is not necessarily absorbing, as recombination may reintroduce a previously absent haplotype. For example, if only *Ab* and *aB* types are found in the population, a recombination event between the two loci may create either an *ab* or *AB* type in an individual in the next generation. Some of the edges of the domain are absorbing, since once one of either *A/a* or *B/b* fixes at the left or right locus, respectively, that two-locus pair remains fixed in the absence of recurrent mutation.

We numerically solved Eq. 2 using finite differencing in a framework similar to Ragsdale et al. (2016). We split the diffusion operator into mixed and non-mixed terms, using an implicit alternating direction scheme for the non-mixed spatial derivatives (Chang and Cooper, 1970) and a standard explicit scheme for the mixed spatial derivatives. We used equal numbers of uniformly spaced grid points for each spatial dimension, so that grid points coincided directly on the off-axes surface of the domain. This allowed for density to be accurately integrated along the surface and interior of the domain. As discussed in Ragsdale et al. (2016) and detailed in the Supporting Information, naively applying finite differencing along the off-axes surface led to numerical error in the solution to *ϕ* Thus, we instead accounted for density moving between the interior of the domain and that surface by directly moving density between the two each timestep.

Because the diffusion equation is linear, it can be used to solve for the density of all two-locus frequencies in the population by allowing for the influx of new mutations each generation. For the single locus diffusion equation, this amounts to the injection of density at rate *θ*/2 at frequency 1/(2*N*), with the appropriate limit taken to allow *N* → *∞*. In the two-locus model, one of the two loci will already be polymorphic (suppose the right *B/b* locus), and a mutation occurs at the other (left) locus. As described above, the new mutation *A* at the left locus initially has frequency 1/(2*N*), while the right locus carries derived allele *B* with frequency x ∈ (0, 1) depending on the single-locus population allele frequency spectrum *f*(*x*), which will itself depend on the population size function *v(τ)*. Allele *A* falls on the *B* background with probability *x* and the *b* background with probability 1 − *x*. Thus, we inject density into the two-locus diffusion equation by simultaneously tracking the single locus allele frequency density function *f* and setting the influx of density into φ proportional to *f* along the *x*_2_ and *x*_3_ axes (Fig. 1B). To solve for the two-locus spectrum under a non equilibrium demographic model *v(τ)*, we first solve for *φ* at equilibrium and then integrate forward according to *v*. We then sample *φ* against the multinomial sampling distribution with sample size *n* (Eq. 1) to obtain the two-locus spectrum.

### Composite likelihood estimation and demographic inference

We follow the composite likelihood approach outlined by Hudson (2001), in which we consider pairs of loci and their sampling distribution. Reducing the full likelihood for more than two linked loci to the composite likelihood over all possible pairs of polymorphisms leads to the loss of information. However, computing two-locus sampling statistics retains a considerable amount of information regarding both allele frequencies and patterns of linkage disequilibrium between them. For recombination distances *ρ* ∈ [*ρ*_min_, *ρ*_max_], we consider all pairs of loci separated by each *ρ*, and store sampling frequencies in the two-locus frequency spectrum for this range or *ρ*. In practice, recombination distances vary continuously over any interval, so we are required to bin our data within subintervals of *ρ* by defining intervals [*ρ*_0_, *ρ*_1_), [*ρ*_1_, *ρ*_2_),…, [*ρ*_*n* − 1_, *ρ*_n_]. For fine enough subintervals, we approximated the expected two-locus spectrum for an interval (*ρ*_*i* − 1_, *ρ*_*i*_) using our diffusion approach with the mean recombination rate over that interval *ρ* = (*ρ*_*i* − 1_ + *ρ*_*i*_)/2.

For a given *ρ*-interval, we made the assumption that all pairs of loci contributing to the two-locus spectrum are independent, approximating the full likelihood by the composite likelihood across all pairs of loci. The two-locus frequency spectrum then forms a Poisson random field, so for sample data *D* and expected model *M* calculated under model parameters Θ, the likelihood of the data 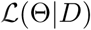 can be calculated by assuming each data entry *D_i_* is a Poisson random variable with mean *M_i_*. Thus, the likelihood function for a single *ρ*-bin is

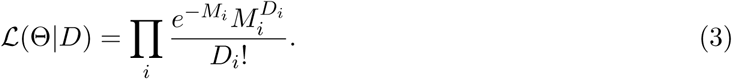

We allowed the population mutation rate *θ* to be an implicit parameter for each bin, which scales the total size of the frequency spectrum while retaining its shape. The maximum likelihood value for *θ* is then 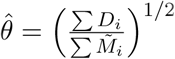, where 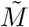 is the model spectrum with *θ* set to one. The square arises because mutations that are paired to existing variant sites arise proportional to rate θ, but those existing mutations also arise proportional to rate *θ*, so that the total rate of influx of new two-locus pairs occur at a rate proportional to *θ*^2^.

We simultaneously considered all bin intervals of *ρ* ∈ [*ρ*_min_, *ρ*_max_], and so for bin centers (*ρ*_1/2_, *ρ*_1 + 1/2_,…), the likelihood function is

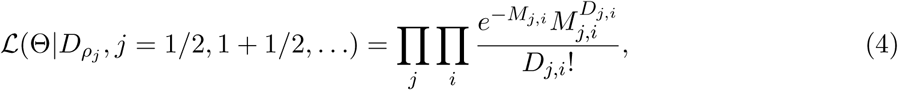
where *j* indexes the *ρ*-bins, and *i* indexes the frequency spectrum entries for a given *ρ_j_*. In reality, pairs of loci are not independent, so we used the Godambe Information Matrix (GIM) to estimate parameter uncertainties (Co man et al., 2016), which adjusts the composite likelihood statistics to account for linkage between data. This required bootstrapping the data, and we did so by dividing the autosomal genome into 1,000 bins of equal length and resampling these regions with replacement.

We fit single-population demographic models to the data, which are defined by the population size history function *v(τ)* (Eq. 2). We considered simplified demographic models that may be described by a handful of parameters, rather than inferring a parameter free function *v(τ)* as in Liu and Fu (2015). For example, in an instantaneous expansion model, the parameters are the relative change in size *v* and the time *T* in the past that the population changed size.

### Phased and unphased data

For data with phased chromosomes, determining haplotype frequencies is straightforward counting of haplotypes for a given pair of loci. Using an aligned outgroup, the ancestral state for each SNP may be determined, so that the two-locus spectrum stores derived two-locus allele frequencies. The ancestral state for each locus may be misidenti ed, potentially due to sequencing error or recurrent mutation along the lineage leading to the outgroup, and this can distort the two-locus spectrum (Hernandez et al., 2007). To account for ancestral misidentification, we included the probability *p*_mis_ ∈ [0, 1] that a given SNP had a misidentified state in our model fitting. Thus, with probability *p*_mis_(1 − *p*_mis_) the *A* allele was misidentified but the *B* allele was correctly identified, and with the same probability the *B* allele was misidentified and the *A* allele was correctly identified. Both alleles *A* and *B* were misidentified with probability 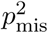. In our demographic model ts to data, we fit *p*_mis_ along with the parameters from the demographic model.

When data is unphased, as is the case for many genomic datasets, observed haplotypes can not be tallied. Rather, we are left with counts of genotypes in individuals, (*n*_AABB_, *n*_AABb_, *n*_AAbb_, *n*_AaBB_,…). The composite linkage disequilibrium statistic 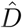 is an unbiased estimator for *D* (Weir, 1979)

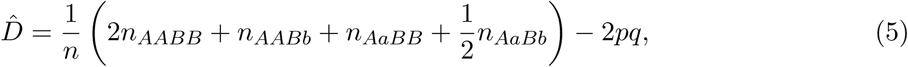
361

Where *n* is the number of sampled individuals. One possible approach to summarize observed data might be to work with the joint statistics *p* = *n_A_*, *q* = *n_B_*, and 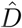. Instead, we directly used genotype counts in the "genotype frequency spectrum" *G*. In genotype data, individuals may carry *AA*, *Aa*, or *aa* at the left locus, and *BB*, *Bb*, or *bb* at the right locus. Thus, there are nine possible two-locus genotypes (*AABB*, *AABb*, *AAbb*, *AaBB*,…) that could be observed to be carried by an individual, so that *G* is an eight dimensional object with size (*n* + 1)^8^. However, *G* is sparse and can be stored efficiently. Each genotype can only be formed by the pairing of two speci c haplotypes (e.g. *AABb* can only be from one haplotype of each *AB* and *Ab*), except for *AaBb*, which could be formed by *AB*+*ab* or *Ab*+*aB*. Thus, we expected *G* to still carry information about demography through the joint patterns of allele frequencies and linkage disequilbrium. Expected genotype frequencies can be calculated from expected haplotype frequencies, and we detail our approach in the Supporting Information.

### *Drosophila* sequence data and recombination map

As an application, we considered a single Zambian population of fruit flies, using data from phase 3 of the *Drosophila* Population Genomics Project (DPGP3), available from the *Drosophila* Genome Nexus (Lack et al., 2015). The data consisted of 197 sequenced haploid embryos, so genomes were necessarily phased. We used Annovar (Wang et al., 2010) to annotate all biallelic SNPs across the genome, and we used intronic and intergenic regions in our two-locus analysis. We determined the ancestral allele for each SNP using the alignment to *D. simulans* (April 2006, dm3 aligned to droSim1, downloaded from the UCSC genome browser), by assuming the *D. simulans* allele was ancestral. If the *D. melanogaster* site had no alignment, or if the *D. simulans* allele was different than the two *melanogaster* alleles, we discarded that site.

For each chromosome, we considered all pairs of biallelic SNPs in intergenic and intronic regions for which an ancestral state could be determined, within recombination distance *ρ*_max_. We determined recombination distances using the recombination map inferred by Comeron et al. (2012), which reports cumulative recombination rates in units of cM over 100,000 bp intervals along each chromosome. We converted to *ρ* = 4*N_e_r* by taking the map distance *d* (in cM) separating the two SNPs and multiplying by 4*N*_e_/100. This required an estimate for *N_e_*, so we used neutral demographic fits to intronic and intergenic single-locus data, which provided an estimate for *θ* = 4*N_e_ μ L*. Here, is the mutation rate, and we used *μ* = 5.5 × 10 ^−9^ (Schrider et al., 2013). The total length of sequences that were included in our analysis was *L* ≈ 3.93 × 10^7^. Then *N_e_* = *θ*/(4μ*L*) ≈ 3 × 10^5^. For each two-locus pair, we counted the number of *AB*, *Ab*, *aB*, and *ab* haplotypes across all 197 samples and then subsampled to a sample size of *n* = 20. In the supporting information, we show how to project data to a smaller sample size, but for the sample sizes in our dataset the full projection would have required more memory than we had available. This allowed for more pairs to be included in the data, as any pair of loci without missing haplotype data for at least 20 samples was included, and a smaller sample size allowed for more rapid evaluation of the expected frequency spectrum for optimization.

### Independent inference of *N_e_*

Two-locus statistics are binned by the populations size-scaled recombination rate *ρ* = 4*N_e_r*, where *r* is the recombination rate per meiotic event per generation. Thus, given a recombination map we require an accurate estimate for *N_e_* to appropriately bin the data. In the case that the effective population size is unknown, *N_e_* may be left as a parameter to be fit during optimization of the model to the data. In this approach, we guess an initial effective population size *N_0_* to first bin the data by *ρ*_0_ = 4*N*_0_*r* (for example, 10^4^ for human populations, or 10^6^ for *Drosophila*) and then allow the *ρ*-value for each bin to be rescaled by *α_N_* as *ρ* = 4*N*_0_*rα_N_*. If the best fit *α_N_* = 1, then *N*_0_ turned out to be the best fit effective population size, while if *α^N^* is larger or smaller than one, then the best fit *N_e_* is inferred to be larger or smaller than *N*_0_ by that factor. We rescaled the ρ value for each bin of data instead of reassigning data to fixed bins for fair comparison of likelihoods across varying values of *α_N_*, and because reassigning two-locus data each iteration of optimization would be computationally burdensome.

## Results and Discussion

### Numerical accuracy of solution to two-locus allele frequency spectrum

We first compared our numerical solution for two-locus statistics for a population in demographic equilibrium to those calculated by Hudson (2001). Our solution matched those using Hudson’s algorithm across all values of *ρ*, from completely linked (*ρ* = 0) to loose linkage (*ρ* = 100) (Fig. 2, top row). To verify our numerical solution for nonequilibrium demography, we compared it to simulations of the discrete two-locus process with an influx of mutations. We simulated a population of *N* = 1000 diploid, randomly mating, individuals for independent pairs of loci separated by a given recombination rate. New two-locus pairs entered the population at a rate proportional to Eqs. S3 and S4. We allowed the simulation to proceed for 20*N* generations and then applied specified population size changes, sampling two-locus haplotype frequencies from the population after each simulation completed. Our nonequilibrium solution matched the simulated two-locus statistics (Fig. 2, bottom row). See Supporting Information for further details regarding simulation and numerical accuracy.

**Figure 2:**
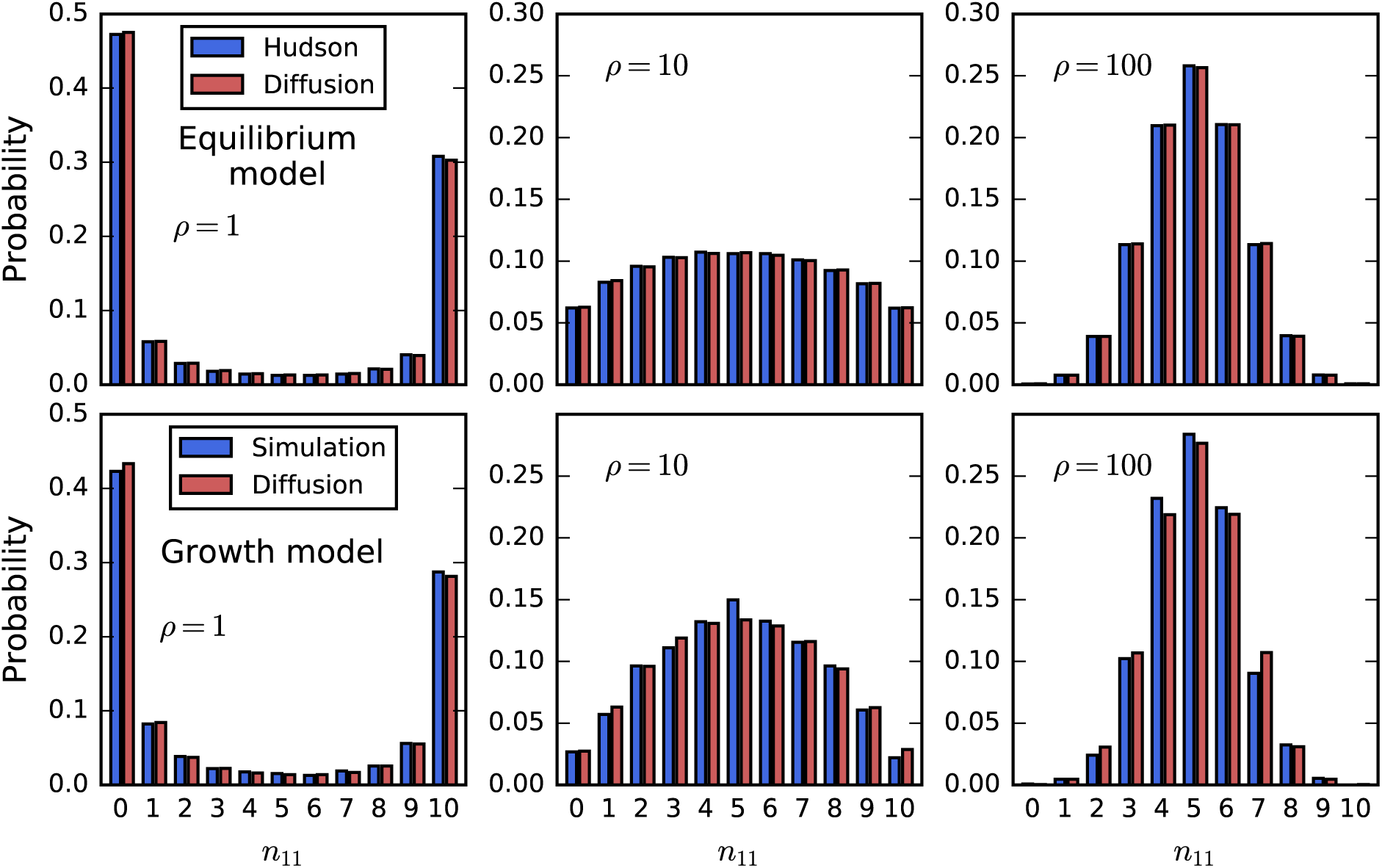
Verification of numerical solution. For sample size *n* = 30, the distribution of *n_AB_* is shown, when the frequencies of *A* and *B* are *p* = 10 and *q* = 15 and *ρ* is varied. Top row: Comparison to equilibrium statistics from Hudson (2001). Bottom row: Comparison to discrete simulation under growth model.

### Two-locus statistics are sensitive to demography

To assess the increase in statistical power for demographic history inference using the two-locus spectrum versus the single-locus spectrum, we used the information theoretical measure Kullback-Leibler (KL) divergence (Kullback and Leibler, 1951). The KL divergence measures the amount of information lost if an incorrect demographic model *M*_0_ is used to approximate the true model *M*_true_, and it can be interpreted as the expected likelihood ratio statistic for testing *M*_true_ against *M*_0_. For discrete distributions, such as frequency spectra, KL divergence is defined as

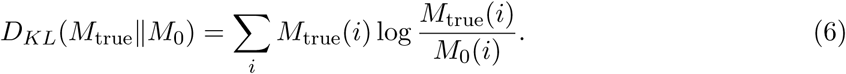

In our comparisons, we took *M*_0_ to be a model of constant demography and compared the KL divergence for two demographic models, an instantaneous growth model and a bottleneck and recovery model, between two-locus and single-locus frequency spectra (Fig. 3). A larger KL divergence indicated that more information is contained in the data to reject the constant size model. For the two model types, we considered varying recovery times *T* since the demographic event, so in the growth model *T* is the time since the instantaneous expansion (*v* = 2), and in the bottleneck model *T* is the time since recovery from the bottleneck (*v_B_* = 0.1; *T_B_* = 0.05). In all cases, the two-locus spectrum is more informative about the demography per pair of linked loci than are two unlinked loci in the single-locus frequency spectrum.

**Figure 3:**
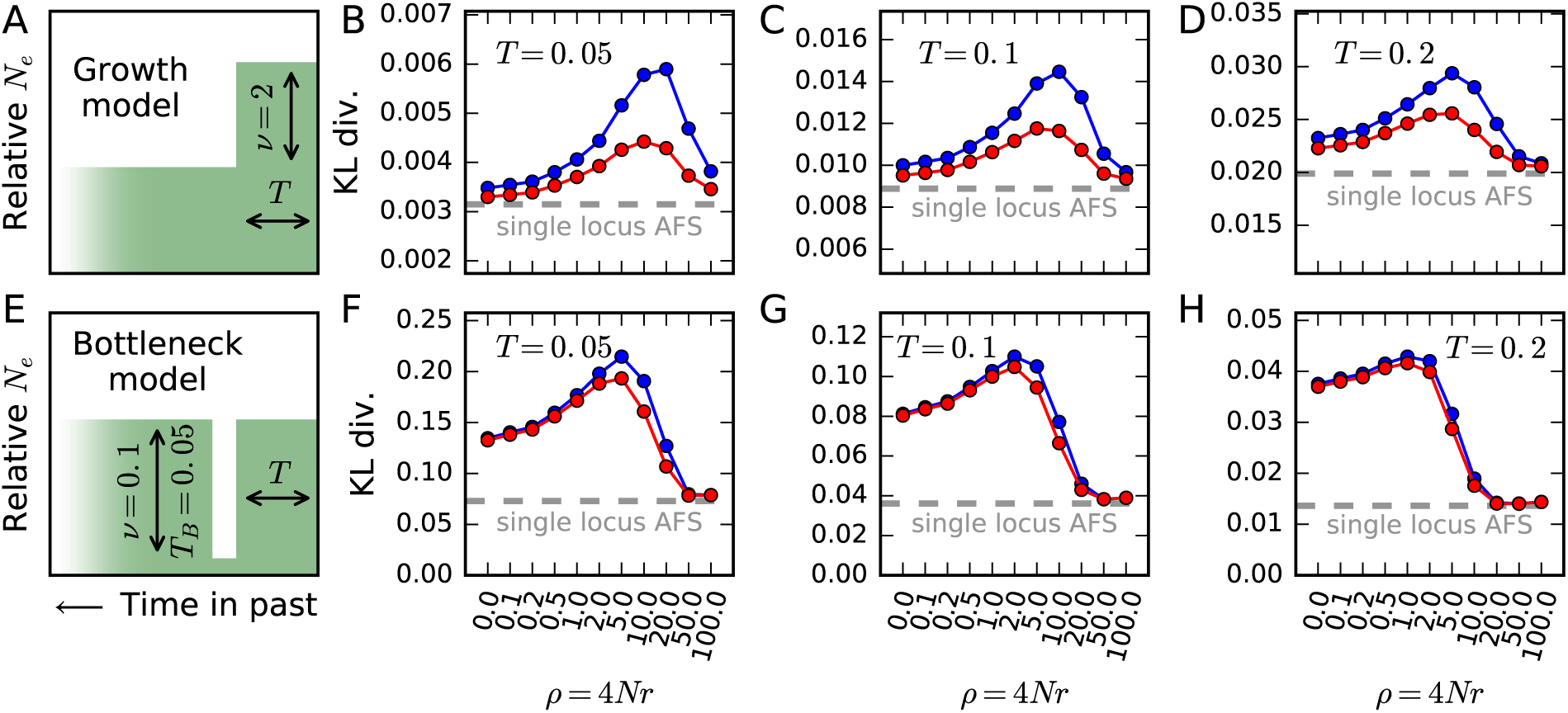
Sensitivity to demography. We compared KL divergence measures between two-locus statistics and the single-locus frequency spectrum for a simple growth model (A, top row) and a bottleneck model (E, bottom row). The blue curve shows the KL divergence for phased (haplotype) data, while the red curve is for unphased (genotype) data. In each comparison, we considered the KL divergence between the specified demographic model and a null model of constant population size. (A) In the instantaneous growth model, the population doubled in size some time *T* in the past, and we considered (B) *T* = 0.05, (C) 0.1, and (D) 0.2. (E) In the bottleneck model, the population shrank to 1=10 its original size for *T_B_* = 0.05 genetic time units and then recovered to its original size *T* genetic units ago for (F) *T* = 0.05, (G) 0.1, and (H) 0.2. In all cases, and across all values of *ρ*, KL divergence was greater for two-locus statistics than the corresponding single locus statistics of the same number of unlinked sites. The two-locus spectrum is thus more sensitive to demographic history than the single-locus spectrum.

We considered the KL divergence for varying values of recombination rate *ρ* from completely linked (*ρ* = 0) to loose linkage (*ρ* = 100). For large *ρ*, KL divergence from two-locus statistics converged to the measure for unlinked single-locus data, which is to be expected as *ρ* → ∞ implies unlinked loci. Importantly, the most informative recombination distance varied between demographic models and recovery times *T* since demographic events. As *T* increases, lower recombination rates are relatively more sensitive, because higher recombination rates will restore levels of linkage disequilibrium faster than lower recombination rates. Therefore, loosely linked loci are more informative about recent demographic events, while tightly linked loci ar emore informative about deeper events. We performed the KL divergence analysis on genotype data as well (Figure 3, red curves), and we found that two-locus statistics at the genotype level are also more sensitive than one-locus statistics. For the growth model, the KL divergence of genotype data was intermediate between the KL divergences of one-locus and haplotype data, but for the bottleneck model, very little sensitivity is lost when using genotype data instead of haplotype data.

### Fits to simulated data

To further validate our model and to explore efficient and informative ways to collate two-locus statistics, we simulated single-population demographic history under neutrality with realistic human mutation and recombination rates for many large (1 Mb) regions using ms (Hudson, 2002) (details in Supporting Information). Using sets of 100 simulated 1 Mb regions, we simulated a simple growth model (instantaneous expansion by a factor of 2, 0:1 time units before present) and the demography to both simulated single- and two-locus statistics (Supporting Information). We repeated this simulation and fitting process 50 times and checked how accurately and precisely we recovered the simulated demographic parameters. We used the same simulations to check the accuracy of our fits to genotype data, by pairing chromosomes to create diploid individuals. Fig. 4 shows our fits to simulated data, with two-locus genotype statistics more precisely recovering the true demographic model than single-locus statistics, and haplotype statistics more precisely than genotype statistics. When we allowed *N_e_* to vary, we also accurately recovered the simulated parameters including *α_N_* (Fig. 4B).

**Figure 4:**
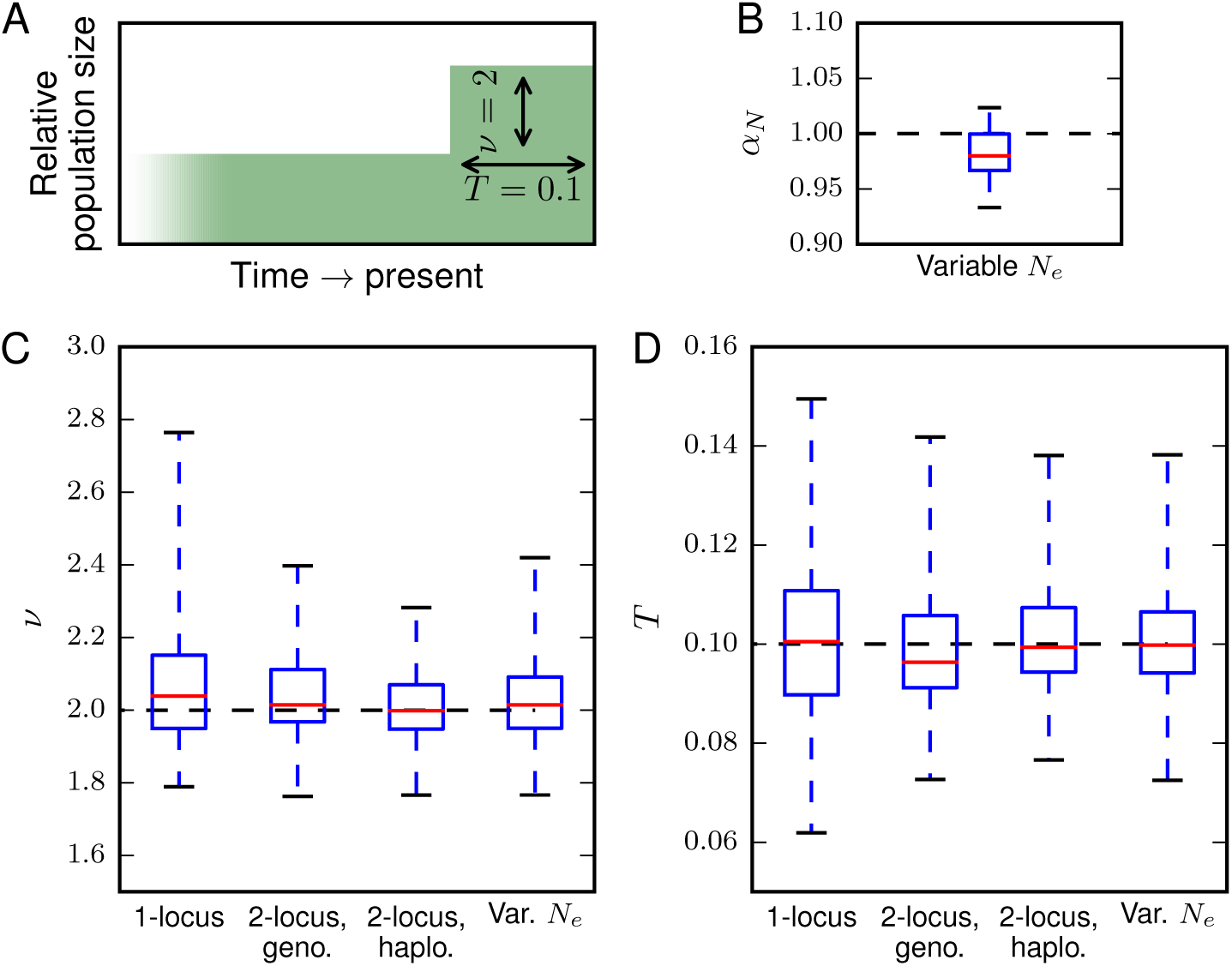
Fits to data from simulated growth model. A: We simulated 50 replicate data sets with length 100Mb under an instantaneous growth model using ms and checked how accurately we recovered the simulated parameters for both single-and two-locus data, including allowing *N_e_* to vary (B). C-D: For both *v* and *T*, ts to the two-locus frequency spectrum were more accurate than single-locus fits. Here, the median values and top and bottom quartiles are indicated by the boxes, and the whiskers extend to the largest and smallest inferred values from the simulated datasets.

In an identical fashion, we also simulated a bottleneck model, in which the population size shrank by a factor of 0.1 for 0.05 genetic time units and then recovered to its original size for 0.2 time units until sampling at present (Fig. 5). For this demography, the fits to single-locus statistics were inconsistent, and many replicates did not converge to reasonable parameter values, with *v_B_* tending to 0. The two-locus haplotypets more accurately recovered the modeled parameters, although the inferred values of *v_B_* were consistently slightly elevated. The fits to genotype data were also more accurate than using single-locus data, consistent with our KL divergence results (Figure 3). Disentangling the depth and duration of a bottleneck from allele frequency data is notoriously challenging (Keinan et al., 2007; Bunnefeld et al., 2015), and jointly incorporating information about linkage disequilibrium dramatically improves parameter identifiability.

**Figure 5:**
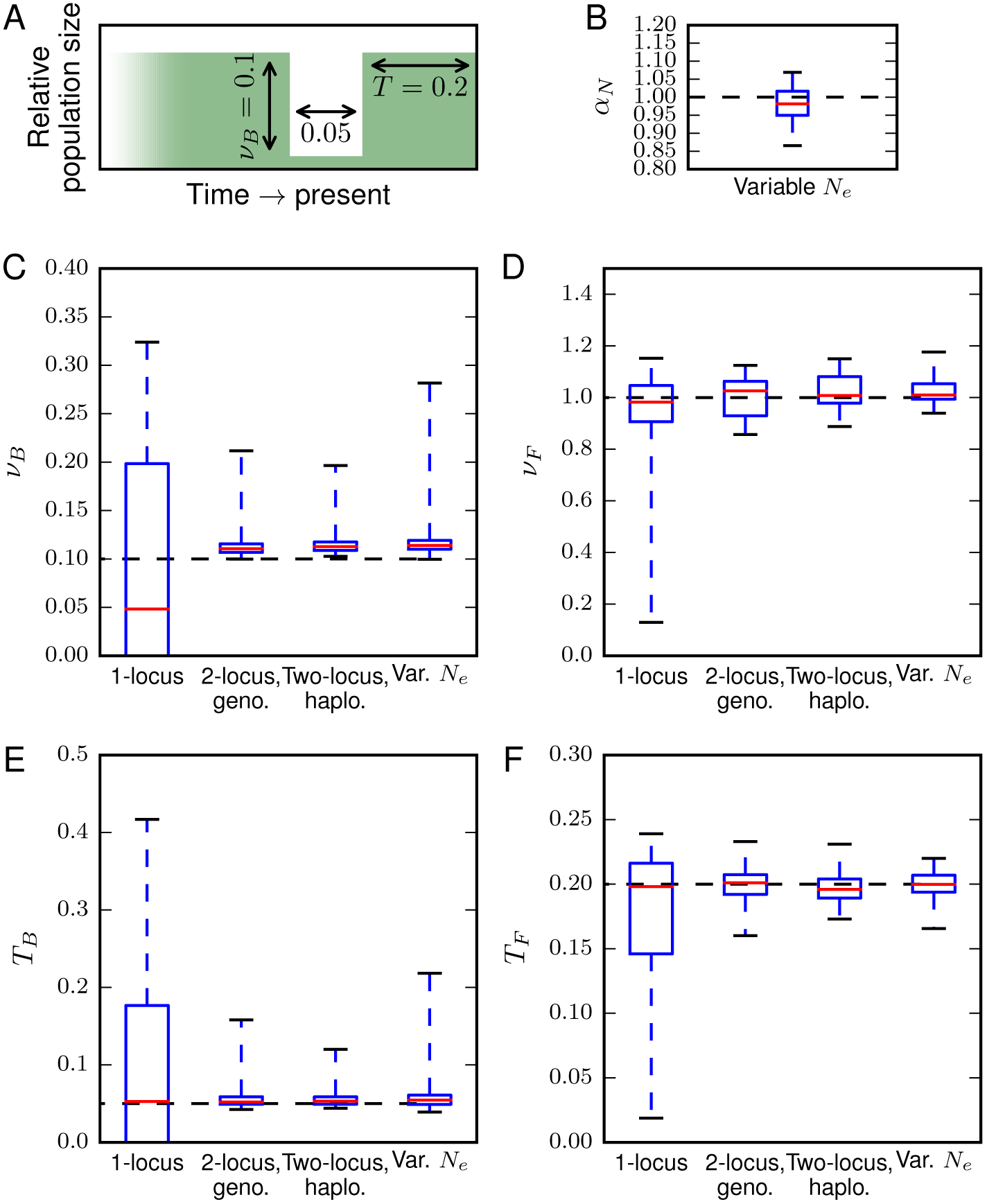
Fits to data from simulated bottleneck model. A: We simulated 50 replicate data sets with length 100Mb under a bottleneck and recovery demographic history, in which the population declined to 0.1 its original size for *T* = 0.05 genetic time units and then recovered to its original size for 0.2 time units. C-F: Demographic inferences using single-locus data alone could not consistently recover the true parameters. However, using genotype or haplotype two-locus data allowed for precise inference of model parameters, including when *N_e_* was allowed to vary (B).

### Demographic inference of a Zambian Drosophila population

As an application of our approach, we considered the demographic history of a Zambian population of *Drosophilia melanogaster*, which is thought to be a close proxy to the ancestral population (Lack et al., 2015). We first fit two- and three-epoch single-population demographic models to intronic and intergenic single-locus data in order to estimate *θ* and *N_e_* (Table 1). We inferred the ancestral effective population size to be approximately 3 × 10^5^, which is somewhat lower than previously suggested sizes for *D. melanogaster* (Keightley et al., 2014; Garud and Petrov, 2016). Using the recombination map of Comeron et al. (2012), we determined distances in *ρ* between pairs of loci, assuming an effective population size of 3 × 10^5^, and we binned two-locus data as described above. We then t the two-and three-epoch models to the two-locus data, with and without varying *N_e_* (Table 1) and calculated parameter uncertainties using the Godambe Information Matrix (Table S1). For all fits, we subsampled the data to 20 samples for computational speed, and additional speed-up was afforded by calculating each *ρ*-bin’s expected frequency spectrum in parallel.

For the two-epoch model, parameter values inferred using single- and two-locus data were quite similar (Table 1). For the three-epoch model, however, inferred values were quite different. In particular, the two-locus fit with fixed *N_e_* inferred a large population size increase followed by a sharp decline, but the single-locus fit and the two-locus fit with variable *N_e_* both inferred two-stage increases with qualitatively similar estimates. When we allowed *N_e_* to be simultaneously fit to the data, we found the best-fit value was smaller (1.7 × 10^5^), and the variable *N_e_* three-epoch model best fit the two-locus data. The disagreement of inferred parameters for the fixed-*N_e_* fit is likely due to the model attempting to fit observed LD but being constrained by an *N_e_* larger than the optimal value. This suggests that scaling the recombination map by a fixed estimate for *N_e_* may introduce significant bias into downstream parameter estimates.

All of the inferred models fit the single-locus frequency spectrum well (Fig. S2), but they varied in their ability to capture patterns of LD (Fig. 6). The two-locus data fit with a three-epoch model including variable *N_e_* fit the LD decay curve much better than any of the other model ts, although it still underestimated long-range LD. Previous models of *D. melanogaster* demographic history also underestimated long-range LD (Garud and Petrov, 2016). While a more complex demography might be able to better fit the LD curve, factors aside from single-population demography may be critical to generating the pattern of long-range elevated LD, including population substructure, recent admixture, or the effects of linked selection.

**Figure 6:**
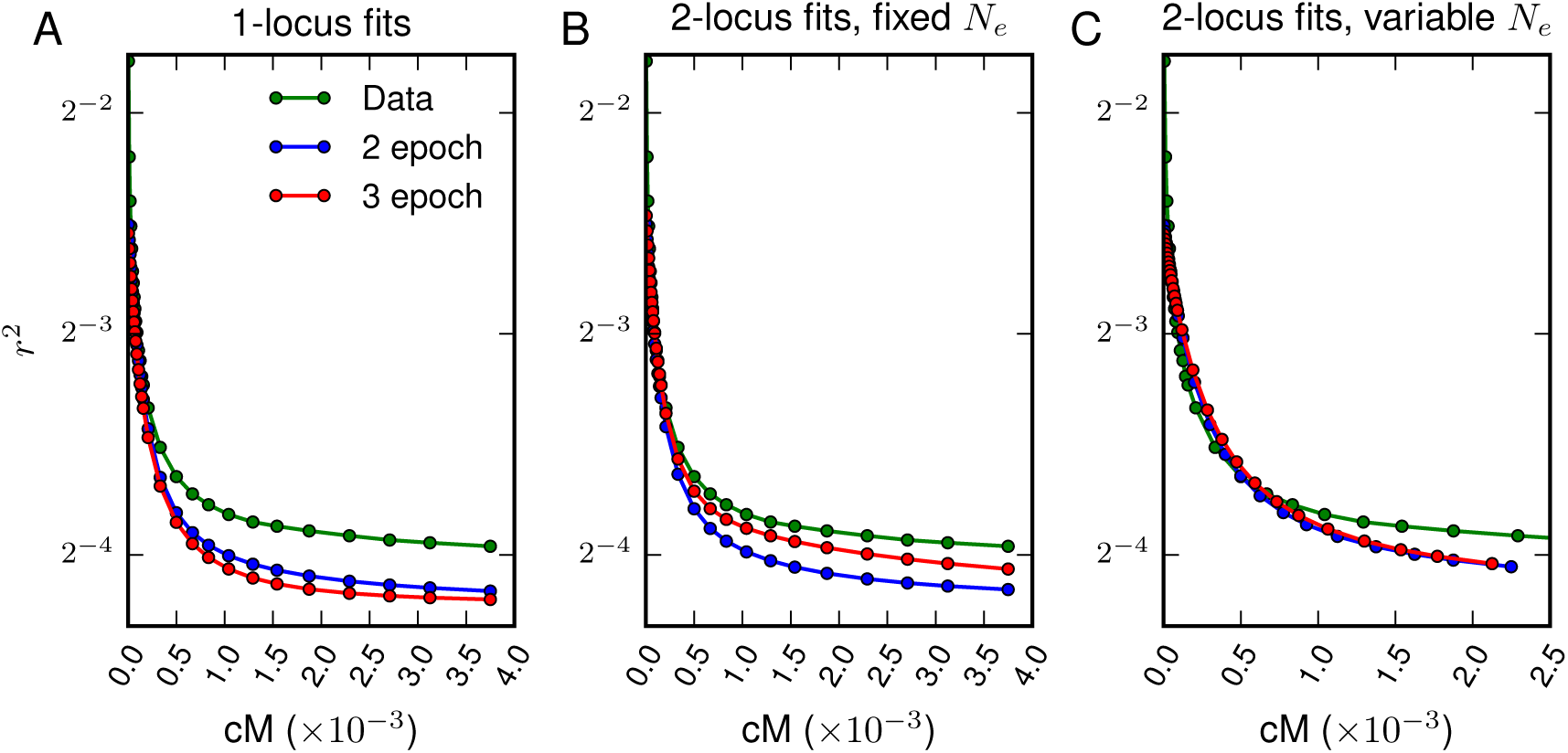
Fits to LD-decay from *Drosophila* data. LD-decay curves for two-locus models compared to observed decay curves from the data. (A) The two-locus model using the best fit parameters from single-locus data, (B) the two-locus model fit with *N_e_* set to 3 × 10^5^, and (C) the two-locus model with *N_e_* allowed to vary. Each of the models underestimates long-range LD decay, as also observed by Garud and Petrov (2016), although the two-locus fits that allow variable *N_e_* attempt to compensate for the poor fit to observed levels of LD (C).

Our estimates of the ancestral effective population size of D. melanogaster are notably smaller than previous estimates. Keightley et al. (2014) estimated the spontaneous mutation rate by sequencing a family of two parents and 12 full-sibling offspring and used their estimation to infer *N_e_* ∼ 1.4 × 10^6^. The effective population size may also be estimated from observed levels of diversity, and Charlesworth (2015) estimated *N_e_* ∼ 0.7 × 10^6^ using observed synonymous site diversity. Furthermore, *N_e_* is often assumed to be ≥ 10^6^ in many population genetic studies of *D. melanogaster* (Sella et al., 2009; Garud et al., 2015; Garud and Petrov, 2016). Our estimates for *N_e_* were substantially lower. Using levels of diversity for intronic and intergenic loci, we estimated *N_e_* ∼ 3 × 10^5^ through our demographic fits to the single-locus AFS (Table 1). In an alternative approach, we allowed *N_e_* to vary in the two-locus inference, and we estimated a smaller value of *N_e_* ∼ 1.7 × 10^5^. This approach is based on the rescaling of the recombination map without assuming a xed mutation rate, and it thus provides an independent inference of the effective population size. Together, our results suggest that ancestral *N_e_* for *D. melanogaster* may be substantially lower than previously estimated, and studies that require an assumed effective population size should consider a wider range of possible *N_e_* values. Notably, it has been suggested that linked selection is common throughout the genome of *D. melanogaster* (Garud and Petrov, 2016), and linked selection is known to increase the variance in offspring distribution, which in turn decreases the effective population size (Le er et al., 2012).

## Conclusions

Based on the continuous approximation to a two-allele two-locus discrete Wright-Fisher model with recombination, we developed a numerical solution to the two-locus diffusion equation that handles arbitrary recombination rates and demographic history. We used this method to develop a composite likelihood framework to infer demographic history from observed two-locus data, which can handle data sampled as either haplotypes or genotypes. While two-locus statistics have been successfully and extensively used to infer fine-scale recombination maps for many organisms, we focused on quantifying the additional power afforded by two-locus over single-locus statistics for demographic history inference. We found that two-locus statistics do provide substantial additional power. For example, while inferring the parameters of a bottleneck model from single-locus data is notoriously difficult (Keinan et al., 2007), we were able to precisely and consistently recover the correct demographic parameters using two-locus statistics. Moreover, for at least some scenarios, little power is lost when data are unphased and genotype frequencies are fit. Finally, we turned to data from a Zambian fruit fly population, and we found that using two-locus statistics to infer demographic history provided a much better fit to both the allele frequency spectrum and observed patterns of LD. The demographic history that we inferred still underestimates the observed long-range levels of LD, which has been previously observed in this population (Garud and Petrov, 2016). Moreover, using two independent approaches, one based on levels of diversity and the other based on scaling the recombination map, we inferred the ancestral effective population size to be substantially lower than previous inferences. It is likely that additional factors to single population demography are at play, including potentially complicated demographic features such as substructure and admixture, and the e ects of linked selection.

## Acknowledgments

AR thanks Gleb Zhelezov for helpful discussions regarding various numerical approaches for this class of PDEs. The authors also thank Nandita Garud for sharing recombination map data and useful discussion regarding the demographic history of the fly population. This work was supported by the National Science Foundation (DEB-1146074 to RG).

## Supporting Information

### Two-locus solution numerics

Our numerical solution to two-locus diffusion equation (Eq. 2) uses finite differences, closely following the numerical methods described in Ragsdale et al. (2016). We separately apply the mixed and non-mixed spatial derivatives, using an alternating direction implicit (ADI) method for non-mixed terms and a standard explicit term for the mixed terms. The grid spacing is uniform with equal number *M* of grid points in each direction *x_i_*, so that grid spacing Δ = 1/(*M* − 1). For the ADI method, each direction was sequentially integrated forward in time. For the *x*_1_ direction, we discretized Eq. 2 as

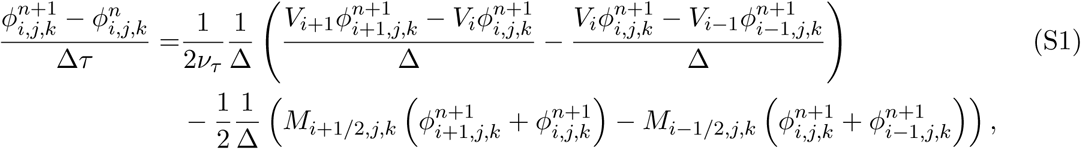
where

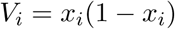
and

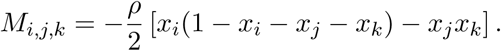

The *x*_2_ and *x*_3_ discretizations were similar, but with the opposite sign for *M_i,j,k_*. For the mixed derivative terms, we sequentially applied an explicit scheme over the (*x*_1_, *x*_2_), (*x*_1_, *x*_3_), and (*x*_2_, *x*_3_) planes. In the (*x*_1_, *x*_2_) direction, we used the discretization

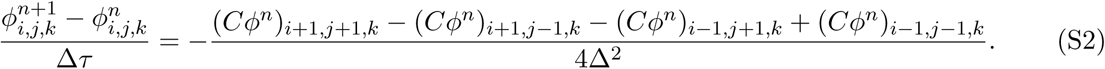

The (*x*_1_, *x*_3_) and (*x*_2_, *x*_3_) planes were analogous.

Sequentially applying the ADI and explicit mixed derivative methods along the o-axes surface resulted in significant error, with an excess of density pushed to the surface. Again, similar to Ragsdale et al. (2016) we integrated *ϕ* forward in time using the methods described above for all grid points not on the o-axes surface. For each grid point near that surface, we calculated the amount of density that should be lost to the surface each time step and directly moved that density to the surface. This density from a grid point at (*x*_1_, *x*_2_, *x*_3_) may be found by numerically integrating the analogous one-dimensional process forward one time unit from a point mass placed at *x* = *x*_1_ + *x*_2_ + *x*_3_ and measuring the amount of density that fixes at *x* = 1. We similarly directly moved density from the surface back into the interior of the domain each time step due to recombination events along that surface. Each time step we also integrated the density on the surface forward in time using Eqs. S1 and S2 for the analogous three state process.

To model the inux of new mutations, we coupled our numerical solution to the two-locus diffusion equation to single-locus models *ϕ^bi^* for the background allele frequencies. These simulations were carried out using 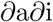 (Gutenkunst et al., 2009), and densities *ϕ^bi^* were added to the two-locus solution *ϕ* along the *x*_2_ and *x*_3_ axes, corresponding to the new haplotype starting at low frequency after mutation. Specifically, suppose *B*/*b* alleles are already segregating at the right locus with the frequency of *B* as *x*, and a new *A* mutation occurs at the left locus. The mutation *A* lands on the *B* background with probability *x* and lands on the *b* background with probability 1 − *x*. We thus added the amount

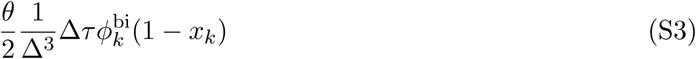
to *ϕ*_0,1,*k*_

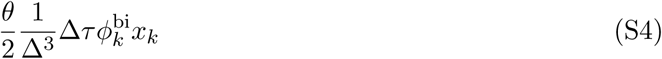
to *ϕ*_0,1,*k*_. The injection for *B* onto *A*/*a* was analogous, adding to *ϕ*_0,*j*,1_ and *ϕ*_1,*j*,0_.

The diffusion equation is valid in the limit of large population size *N_e_*, so we extrapolated on grid spacing Δ to approximate the solutions for Δ → 0. In practice, the number of grid points should exceed the number of samples in the frequency spectrum. With a sample size of 20, we typically used grid spacings with *M* = 40, 50, and 60. We also found that accuracy was improved by extrapolating on Δ*τ* as well, and we used Δ*τ* = [0.005, 0.0025, 0.001] for these grid spacings.

### Binning data by *ρ*

Differences in the two-locus frequency spectra for varying values of *ρ* are more pronounced at small *ρ*. (For example, the differences between spectra for *ρ* = 1 and 2 are much more pronounced than the differences between spectra for *ρ* = 49 and 50.) Thus, we used tighter bins for low recombination rates and wider bins for higher recombination rates. We partitioned data into 28 bins, chosen to match the number of cores on a node of our compute cluster, and computation of spectra for each bin was parallelized. The bin edges were *ρ* = 0, 0.1, 0.2, 0.3, 0.4, 0.5, 0.6, 0.7, 0.8, 0.9, 1, 1.2, 1.4, 1.6, 1.8, 2, 3, 5, 7, 9, 11, 14, 17, 20, 25, 30, 35, 40, and 50.

### Details of simulation using ms

We simulated two demographic models using ms: a growth model and bottleneck model, as described in the Results and Discussion. Each simulation consisted of 100 1Mb regions, and we repeated each simulation 50 times, with a sample size of 20 chromosomes. For both demographies, we set the per-base recombination rate to *r* = 2.5 × 10 ^−8^ and the mutation rate to *μ* = 2.5 × 10^−8^. The input command for the growth model was

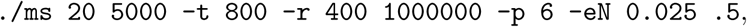
and for the bottleneck model was

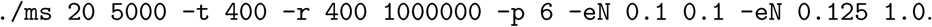

## Projection

In many genomic data sets, some SNPs might not be called in every individual. Moreover, SNPs will vary in the number of individuals for which data exists. Instead of discarding those SNPs with missing data, by projecting the frequency spectrum down to a smaller sample size *n*_proj_, all data called in at least *n*_proj_ sampled chromosomes may be included (Marth et al., 2004). To project the single-locus frequency spectrum from a sample size of *n* to a smaller sample size of *n*_proj_, one averages over all possible ways of picking subsamples of size *n*_proj_ from the *n* observed samples using the hypergeometric function (Marth et al., 2004).

For two-locus statistics, we only included data when both the left and right alleles were called in an individual. To project from *n* observed samples to *n*_proj_, with *n*_proj_ < *n*, we averaged over all possible ways of subsampling the *n* observed haplotypes. For data with sampled haplotype counts 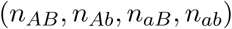,
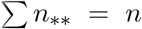, we counted the number of ways to sample 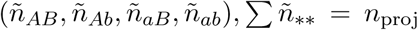 from that collection of *n* samples. The probability that we choose 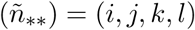 haplotypes from (*n_**_*) can be expressed as

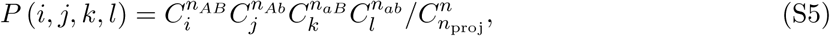

where 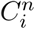 indicates the binomial coefficient with parameters n and i.

### Genotype frequency expectations from haplotype frequencies

For a given entry (*i, j, k*) in the two-locus spectrum with haplotype frequencies

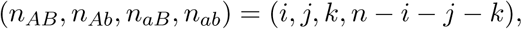

We determined expected genotype frequencies by counting all possible ways that the haplotypes could be paired. To calculate pairing probabilities and visualize the computation, consider pairing a collection of *n* (even) colored balls that could be any of four colors (red, green, blue, and yellow), where *n_R_* is the number of red balls, *n_G_* the number of green, and so forth. The total number of ways than *n* objects can be paired is

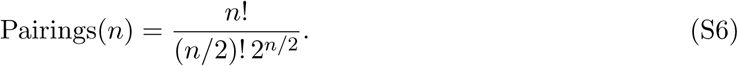

For a given configuration 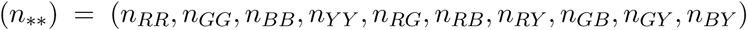, we must also count the total number of ways that the colored balls may be distributed. Here, *n_RR_* is the number of pure red ball pairings in the set, *n_GY_* is the number of pairs of a green and yellow ball paired together, and so forth. First, for pure-colored (e.g. red) pairings, there are 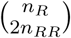 ways to assign red balls between pure and mixed pairings. Of the pure pairings, there are Pairings(2*n_RR_*) (Eq. S6) ways to split the pure red balls into pairs.(The other three colors follow the same calculations.) 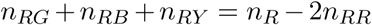 red balls will be paired with non-red balls. For these red balls in mixed pairings, there are 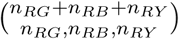 ways to split them into the given number of *RG*, *RB*, and *RY* pairs, where 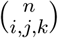 is the trinomial coefficient, with *i* + *j* + *k* = *n*, defined as 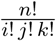. Finally, for red balls that will be paired with green balls, there are n_RG_ permutations of these possible pairings. Again, the other colors follow the same calculation.

Now, the probability that haplotypes with frequencies (*n_R_*, *n_G_*, *n_B_*, *n_Y_*) will the paired as (*n_**_*) is the number of ways that unique pairings lead to that configuration of genotypes, divided by the total number of possible pairings:

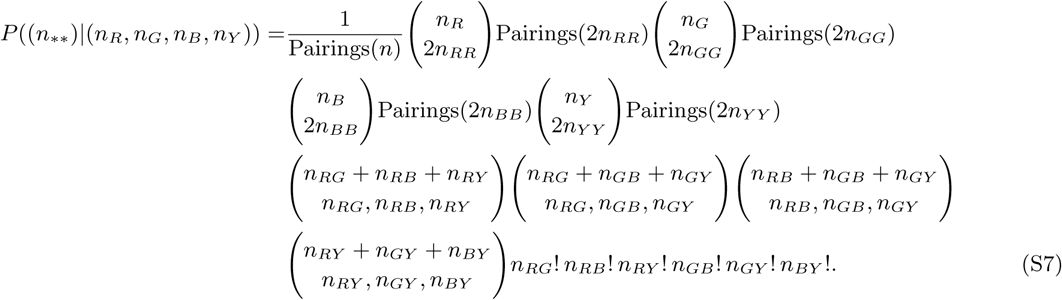

**Table S1:** 95% confidence intervals from ts to *Drosophila* data. We used the Godambe Information Matrix to estimate uncertainties for our best fit parameter values.

| Data (Model) | $\nu_1$ | $\nu_2$ | $T_1$ | $T_2$ | $p_{\text{mis}}$ | $N_e$ |
| --- | --- | --- | --- | --- | --- | --- |
| 1-loc (2-ep) | 4.16 – 4.30 |  | 0.321 – 0.337 |  | 0.0468 – 0.0486 | 295,600 – 310,700 |
| 1-loc (3-ep) | 2.26 – 2.44 | 8.2 – 13.2 | 0.374 – 0.402 | 0.084 – 0.107 | 0.0488 – 0.0504 | 284,500 – 299,000 |
| 2-loc (fix $N_e$, 2-ep) | 3.69 – 3.96 | | 0.358 – 0.383 | | 0.0437 – 0.0460 | |
| 2-loc (fix $N_e$, 3-ep) | 9.03 – 59.6 | 1.64 – 1.75 | 0.209 – 0.231 | 0.0524 – 0.0536 | 0.0422 – 0.0446 | |
| 2-loc (var $N_e$, 2-ep) | 3.94 – 4.10 | | 0.370 – 0.388 | | 0.0450 – 0.0462 | 179,500 – 180,500 |
| 2-loc (var $N_e$, 3-ep) | 1.30 – 1.76 | 4.37 – 4.79 | 0.347 – 0.357 | 0.242 – 0.330 | 0.0460 – 0.0486 | 169,000 – 171,000 |

**Figure S1:**
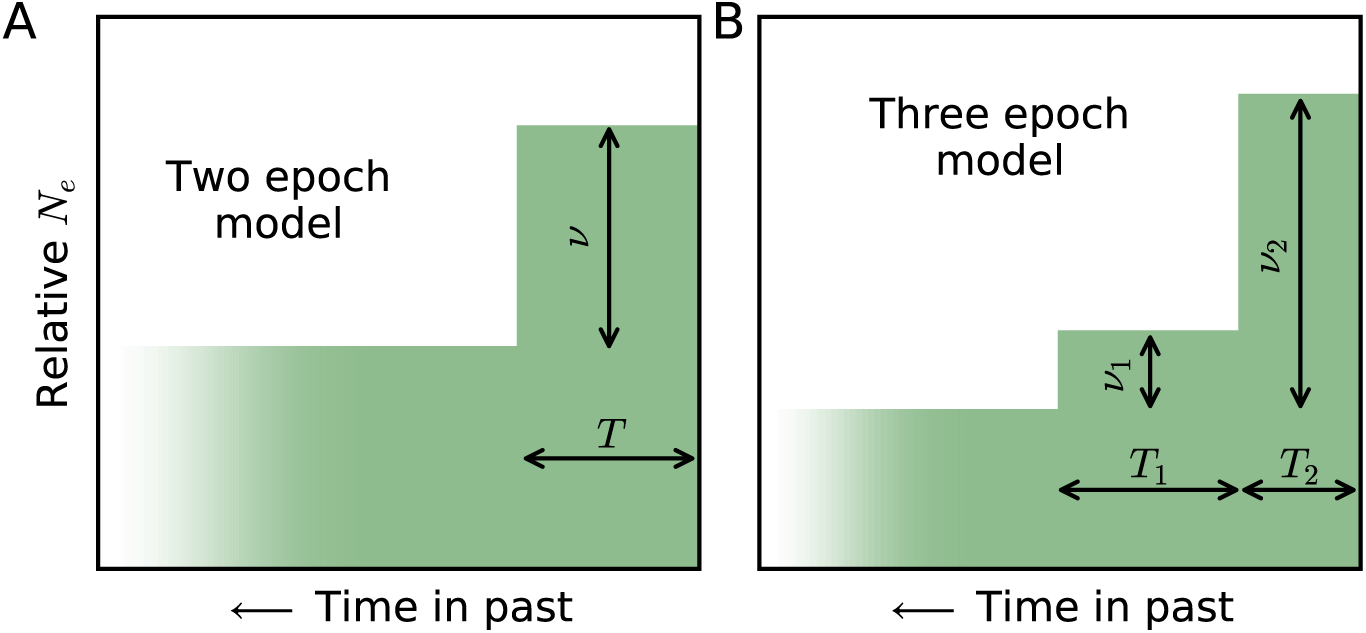
Demographic models fit to data. The two single-population models we simulated data under and then fit to the observed *D. melanogaster* data. (A) The two epoch model has a relative size change *v* some time *T* in the past, while (B) the three epoch model includes two periods of recent size change with sizes *v*_1_ and *v*_2_ relative to the ancestral population size and lasting for times *T*_1_ and *T*_2_, resp.

**Figure S2:**
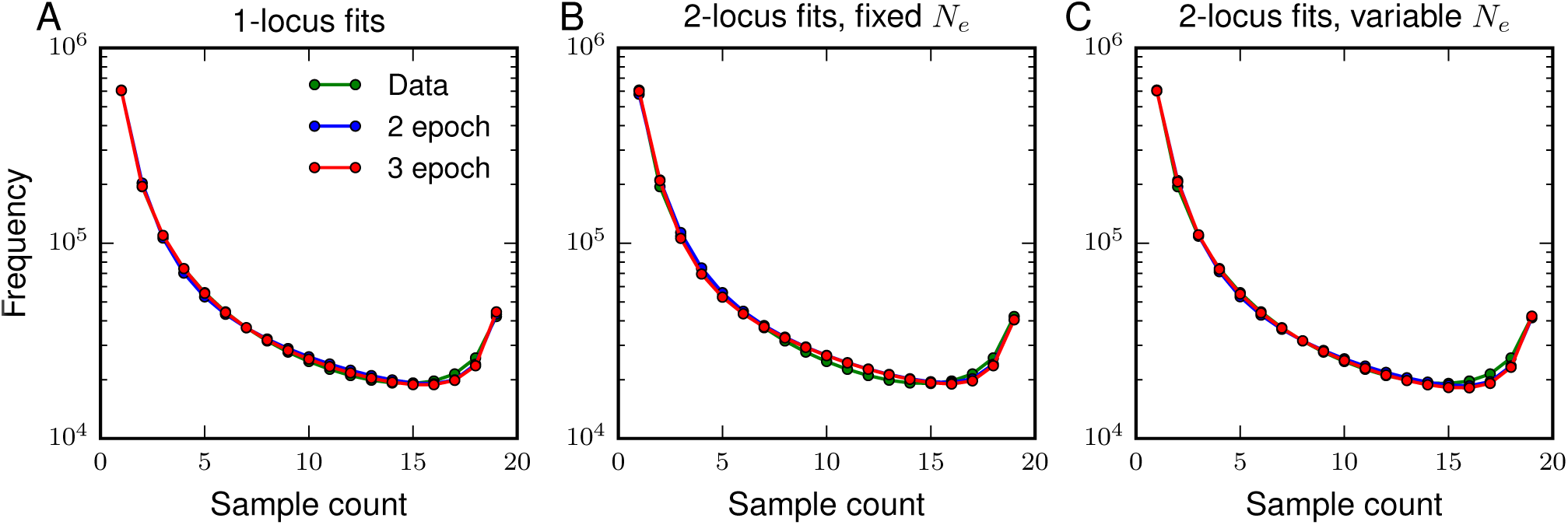
Fits to single-locus AFS. All inferred models fit the single-locus data well. (A) We fit two- and three-epoch models to the single-locus AFS, including a parameter to account for ancestral misidentification that causes the over-representation of high frequency alleles. (B) We fit those same models to two-locus data and fixed *N_e_* = 3 × 10^5^, which was inferred from our ts to the single-locus data. (C) *N_e_* was allowed to vary, rescaling the effective recombination rates.

